# Positive modulation of cerebellar α6GABA_A_ receptors for treating essential tremor: a proof-of-concept study in harmaline-treated mice

**DOI:** 10.1101/2021.04.19.440397

**Authors:** Ya-Hsien Huang, Ming Tatt Lee, Werner Sieghart, Daniel E. Knutson, Laurin R. Wimmer, Dishary Sharmin, James Cook, Marko D. Mihovilovic, Lih-Chu Chiou

## Abstract

**Background:** The etiology of essential tremor (ET) remains unclear but may involve abnormal firing of Purkinje cells, which receive excitatory inputs from granule cells in the cerebellum. Since α6 subunit-containing GABA_A_ receptors (α6GABA_A_Rs) are abundantly expressed in granule cells, we validated a hypothesis that α6GABA_A_R-selective positive allosteric modulators (PAMs) are promising pharmacological interventions for ET therapy.

**Methods:** Employing the harmaline-induced ET model in male ICR mice, we evaluated the possible anti-tremor effects of four α6GABA_A_R-selective PAMs, the pyrazoloquinolinones Compound 6 and LAU-463 and their respective deuterated derivatives. Propranolol, a clinical anti-tremor agent, was employed as positive control. To investigate the involvement of cerebellar α6GABA_A_Rs in the anti-tremor effect of intraperitoneal (*i.p.*) Compound 6, furosemide, an α6GABA_A_R antagonist, was intracerebellarly (*i.cb.*) co-administered with Compound 6. The burrowing activity, an indicator of wellbeing in rodents, was measured concurrently.

**Results:** Harmaline (10-30 mg/kg, *s.c.*) induced action tremor in ICR mice dose-dependently and markedly reduced their burrowing activity. Compound 6 (3 and 10 mg/kg, *i.p.*) significantly attenuated harmaline (20 mg/kg)-induced action tremor and burrowing activity impairment. Propranolol (20 mg/kg, *i.p.*) diminished tremor but failed to restore the burrowing activity in harmaline-treated mice. Importantly, both anti-tremor and burrowing activity restorative effects of Compound 6 (10 mg/kg, *i.p.*) was significantly reversed by co-administration of *i.cb.* furosemide at a dose (10 nmol/0.5 μl) having no effect *per se*. All four α6GABA_A_R PAMs exhibited a similar therapeutic efficacy.

**Conclusion:** α6GABA_A_R-selective PAMs significantly attenuated action tremor and restored physical well-being in a mouse model mimicking ET by acting in the cerebellum. Thus, α6GABA_A_R-selective PAMs may be potential therapeutic agents for ET.

## Introduction

Essential tremor (ET) is a neurological disorder with symptoms characterized by uncontrollable rhythmic shaking of one or more body parts (Deuschl *et al.*, 1998; Haubenberger and Hallett, 2018). It is one of the most common movement disorders, especially in the elderlys (Louis and Ferreira, 2010; Sepúlveda Soto and Fasano, 2020), and a potential risk factor for other neuropsychiatric conditions, such as depression and anxiety (Huang *et al.*, 2019; Louis, 2010). The socio-economic burden inflicted by ET is insurmountable, as it negatively affects the well-being and productivity of the patients and their caregivers (Monin *et al.*, 2017). Current therapeutic agents for ET are limited and often hampered by either inadequate efficacy or intolerable side effects (Shanker, 2019). Therefore, it is an urgent need to develop novel anti-tremor agents with a promising efficacy and minimal unwanted side effects.

The pathogenesis of ET remains not fully understood. Several human studies suggested a general hypo-function of GABAergic transmission in the brains of ET patients. ET patients had lower GABA, but higher glutamate concentrations in the cerebrospinal fluid (CSF) than normal controls (Mally *et al.*, 1996). Positron emission tomography (PET) studies demonstrated that ET patients had a higher binding potential of ^11^C-flumazenil at the benzodiazepine (BDZ) site of GABA_A_Rs than normal subjects in brain regions involved in the pathogenesis of ET, including the deep cerebellar nuclei (DCN), cerebellar vermis, and ventral intermediate nucleus of the thalamus, and these changes correlated with their ET severity (Boecker *et al.*, 2010; Gironell *et al.*, 2012). An increased binding of ^11^C-flumazenil suggests an increased expression of GABA_A_ receptors that contain BDZ binding sites in ET patients and might be a compensatory change in these brain regions due to the reduced GABA levels.

Consistent with a reduced GABAergic inhibition, ET patients have been reported to have excessive cerebellar activity revealed by neuroimaging studies (Sharifi *et al.*, 2014). Postmortem studies in ET patients revealed a significant loss of cerebellar Purkinje cells (PCs) (Louis *et al.*, 2013; Louis *et al.*, 2007) and their dendritic arborization (Kuo *et al.*, 2017; Louis *et al.*, 2007; Louis *et al.*, 2014). Other abnormalities in the cerebellum affected basket cells and climbing fibers (Louis and Faust, 2020). Such changes could have been the cause or the consequence of the excessive cerebellar activity.

In any case, Purkinje cells (PCs) are at the center of the repeating microcircuits that form the cerebellar cortex. They integrate information from five major classes of excitatory and inhibitory interneurons. The cerebellar cortex has two excitatory inputs, the mossy fibers and the climbing fibers. Mossy fibers provide the sensory and contextual information to the excitatory granule cells that elicit the coordinated motor programs via PCs. Climbing fibers provide error signals directly to the PCs, leading to an improved motor program and final timing for motor activity (D’Angelo *et al.*, 2013). Mossy fibers, in addition to granule cells, also excite the inhibitory Golgi cells. Golgi cells directly and indirectly inhibit granule cell excitation via feed forward and feedback loops, respectively, and sharpen their excitatory signal (Mapelli *et al.*, 2014; Nieus *et al.*, 2014). In addition, Golgi cells cause a broad lateral inhibition of granule cells that generates dense clusters of granule cell activity organized in center-surround structures, implementing combinatorial operations of multiple mossy fiber inputs (D’Angelo *et al.*, 2013). The axons of granule cells are ascending to the molecular layer where they originate the parallel fibers that form excitatory synapses on the inhibitory PCs, Golgi cells, basket cells and stellate cells. In contrast to the granule cell-inhibiting Golgi cells, basket cells and stellate cells inhibit PCs, thus increasing the precision of PC firing in response to mossy fiber and climbing fiber excitation (Person and Raman, 2012). PCs are the major output neurons of the cerebellar cortex, providing a precise inhibitory timing control on the neuronal excitability in the downstream deep cerebellar nuclei (DCN) (Hirano, 2019; Pugh and Raman, 2009).

In agreement with their central role in the function of the cerebellum, PCs were recently reported to be significantly involved in the excessive cerebellar activity and play an important role in the pathogenesis of ET. In an ET model in which mice were treated with harmaline, a β-carboline neurotoxin (Lavita *et al.*, 2016), it was demonstrated that the onset of tremor coincides with abnormal rhythmic firing of cerebellar PCs, which alters the downstream firing regularity of DCN neurons, and that a genetic reduction of PC-DCN transmission reduces tremor activity (Brown *et al.*, 2020). This conclusion was also supported by another study indicating that misfiring of cerebellar PCs elicited by synaptic pruning deficits of climbing fiber-to-PC synapses caused by PC-specific glutamate receptor subunit insufficiency, cause excessive cerebellar oscillations and might be responsible for tremor generation (Pan *et al.*, 2020).

Therefore, an enhancement of GABAergic transmission by positive allosteric modulators (PAMs) of GABA_A_Rs, like benzodiazepines (BDZs), may be a therapeutic strategy for ET patients. BDZs indeed are used clinically as a second-line treatment for ET (Shanker, 2019). However, the use of BDZs in ET treatment is limited by their unwanted side effects, like sedation, muscle weakness, addiction, confusion and amnesia (Bruno *et al.*, 2015). Conversely, PAMs that selectively modulate the GABA_A_Rs in the cerebellum would be an ideal option for ET treatment.

GABA_A_Rs are ligand-gated chloride channels consisting of 5 subunits. A total of 6α, 3β, 3γ, δ, ∊, π, θ and 3ρ subunits have been identified in the mammalian nervous system (Olsen and Sieghart, 2008). The majority of GABA_A_Rs consists of two α, two β and one γ or δ subunits (Sieghart and Savic, 2018). The α6 subunit-containing GABA_A_ receptors (α6GABA_A_Rs) are ideal targets for modulating the activity of PCs and a substantial diversity of heterocyclic scaffolds have been employed as allosteric modulators (Vega Alanis *et al.*, 2020). They are insensitive to classical benzodiazepines, like diazepam, (Luddens *et al.*, 1990; Sieghart, 1995) and are massively expressed in cerebellar granule cells (Gutierrez et al., 1996; Pirker et al., 2000). They form α6ßγ2 and α1α6βγ2 GABA_A_Rs (Jechlinger *et al.*, 1998), mediating GABAergic inhibition at the Golgi cell-granule cell synapses (Nusser *et al.*, 1998). In addition, they form α6βδ and α1α6βδ GABA_A_Rs (Jechlinger *et al.*, 1998; Olsen and Sieghart, 2008; Scholze *et al.*, 2020) that are present at extrasynaptic sites at the Golgi cell-granule cell synapse and all over granule cells, their cell bodies, axons and their parallel fibers, mediating tonic inhibition of granule cells (Lee *et al.*, 2010; Nusser *et al.*, 1998). These α6GABA_A_Rs are much less expressed in other brain regions (Gutierrez *et al.*, 1996). Since α6GABA_A_Rs regulate the excitability of granule cells, which represent one of the two main excitatory inputs of the cerebellum, enhancement of their inhibitory efficacy by α6GABA_A_R-selective PAMs seems to be one possible way to reduce the excessive activity of PCs. Here, we confirmed this concept using the harmaline-induced tremor model and four pyrazoloquinolinone compounds (PQs) (Zhang *et al.*, 1995) that we have previously identified to be PAMs highly selective to α6GABA_A_Rs (Knutson *et al.*, 2018; Varagic *et al.*, 2013).

Harmaline can induce action tremor, involving the limbs, tail, trunk, head, and whiskers-, with a frequency range, 10-16 Hz in mice (Martin *et al.*, 2005), 8-12 Hz in rats (Martin *et al.*, 2005), 8-12 Hz in cats (Lamarre and Mercier, 1971) and 6-8 Hz in monkeys (Ohye *et al.*, 1970), resembling the action tremor in ET patients at the frequency of 4-12 Hz (Kuo *et al.*, 2019). Thus, the harmaline-induced tremor model is recognized as an animal model for screening potential therapeutic agents for ET treatment (Martin *et al.*, 2005). The PQ compounds tested include Compound 6, originally coded as PZ-II-029 (**Fig. 1A**), and its structural analogue LAU463 (**Fig. 1B**), as well as their respective deuterated analogues, DK-I-56-1 (**Fig. 1C**) and DK-I-58-1 (**Fig. 1D**), that have a better pharmacokinetic profile with longer half-lives (Knutson *et al.*, 2018). We first established the dose- and time-dependence of harmaline-induced tremor, as well as that of harmaline-induced disruption in the burrowing activity of mice, which is one of the “Activities of Daily Living” in laboratory rodents and has been employed as an indicator of well-being in rodents (Jirkof, 2014). Then, we investigated whether the PQ Compound 6 would inhibit harmaline-induced tremor and restore the burrowing activity disrupted by harmaline. To identify the site-of-action of Compound 6, we subsequently examined whether intracerebellar (*i.cb.*) microinjection of furosemide, a selective α6GABA_A_R antagonist, would reverse the effects of Compound 6. Lastly, we compared effects of the four PQ compounds on tremor and burrowing activities in harmaline-treated mice.

**Fig 1.**
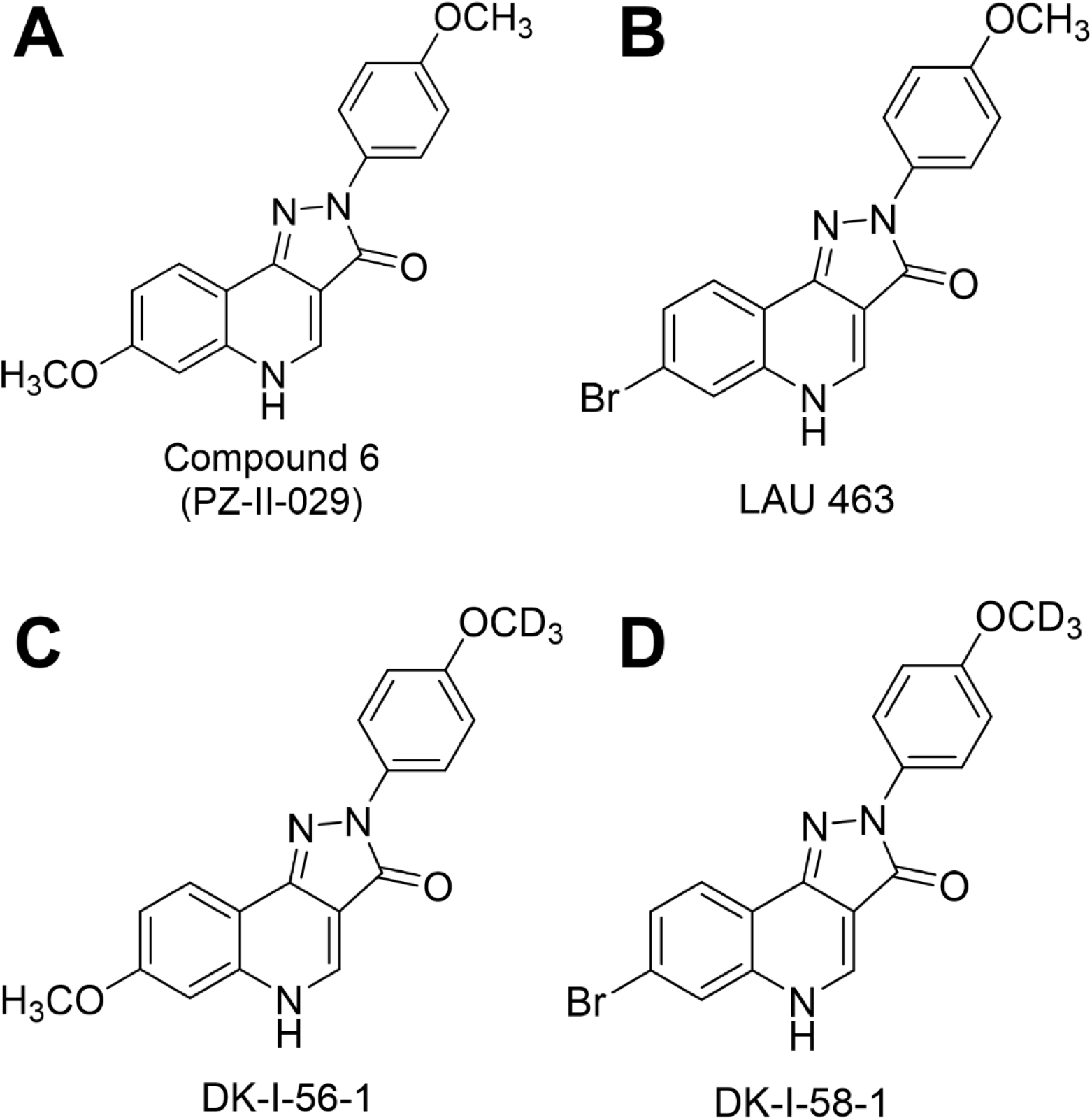
Chemical structures of pyrazoloquinolinone compounds, the α6 subunit-containing GABA_A_ receptor (α6GABA_A_R)-selective positive allosteric modulators (PAMs). (**A**) Compound 6, originally encoded as PZ-II-029, was the first synthesized α6GABA_A_R-selective PAMs. (**B**) LAU463 is a structural analogue of Compound 6. (**C**) DK-I-56-1 and (**D**) DK-I-58-1 are deuterated derivatives of Compound 6 and LAU463, respectively.

## Materials and Methods

### Animals

The animal care and experimental procedures reported in this study were approved by the Institutional Animal Care and Use Committee of National Taiwan University, College of Medicine. Male adult mice (ICR strain) were purchased from BioLASCO (Taiwan Co., Ltd) were housed in groups of 5 in a cage in a holding room with a 12 h light-dark reversed cycle (lights on at 20:00 and off at 08:00) and access to food and water *ad libitum*. On the experimental day, mice (6-10 weeks) were moved with their home cages to a behavior room and acclimated there for at least one hour before testing.

### *Intra-cerebellar (i.cb), intraperitoneal* (*i.p.*) and subcutaneous (*s.c.*) *injections*

The *i.cb.* cannulation procedure was performed as reported in our previous studies (Chiou *et al.*, 2018; Liao *et al.*, 2016) with modifications. Briefly, mice were anaesthetized with sodium pentobarbital (60 mg/kg, *i.p.*) and placed in a stereotaxic frame keeping the bregma-lambda axis horizontal. After shaving the hair and exposing the skull surface, each mouse was implanted with a 24-gauge stainless-steel guide cannula directing toward the vermis (−7.0 mm caudal, −0.4 mm ventral from bregma) according to the stereotaxic coordinate of the mouse (Franklin and Paxinos, 1997). Each mouse was allowed to recover for at least 1 week. On the day for behavioral tests, a 30-gauge injection cannula connected to a 1-μl Hamilton syringe via a 50 cm polyethylene tube was inserted into the guide cannula for drug injection. The drug solution of 0.5 μl was slowly infused with a microinfusion pump (KDS311, KD Scientific Inc.) for 3 min with a further “hold” time for 2 min, while the mouse was allowed to freely move in an open field arena. The microinjection site was confirmed by the positive staining of trypan blue, which was injected through the cannula after the behavioral tests. Data from mice with offsite injections were discarded. For *i.p.* and *s.c.* injection, the drug solution was injected at a volume of 10 ml/kg.

### Tremor induction and measurement

Action tremor was induced in mice by harmaline in accordance with the protocol described previously with minor modifications (Handforth *et al.*, 2018; Martin *et al.*, 2005). The frequency and intensity of tremor were measured and analyzed using a Tremor Monitor (San Diego Instruments, SDI, CA, USA) as reported previously (White *et al.*, 2016), which can accurately differentiate tremor activity from ambulatory/stereotyped movements and global motion activity (Paterson *et al.*, 2009). Each mouse was placed in the chamber for acclimatization, and then the baseline motion power was recorded for 10 min. Tested drugs were administered by *i.p.* or *i.cb.* injection 5 min before harmaline injection. Each raw trace of the movement activity recorded by Tremor Monitor was converted by a fast Fourier transform to generate a frequency domain-based motion power spectrum with the bin size at 1 Hz. The tremor activity in each mouse was measured as the ratio of the motion power at 10-16 Hz over the tremor frequency (0-34 Hz) of the global motion power over a 10-min duration (**Fig. 2A**). The overall tremor activity was expressed by the area under curve (AUC) derived from time-dependent tremor responses. To ensure that the harmaline-induced tremor detected in our setups was indeed action tremor, we videotaped the harmaline-treated mice’s locomotor activity in the tremor chamber and subsequently analyzed the action/immobile motion of the mice using the SMART Video Tracking System (Harvard Apparatus, MA, USA) (Fig.2B).

**Fig 2.**
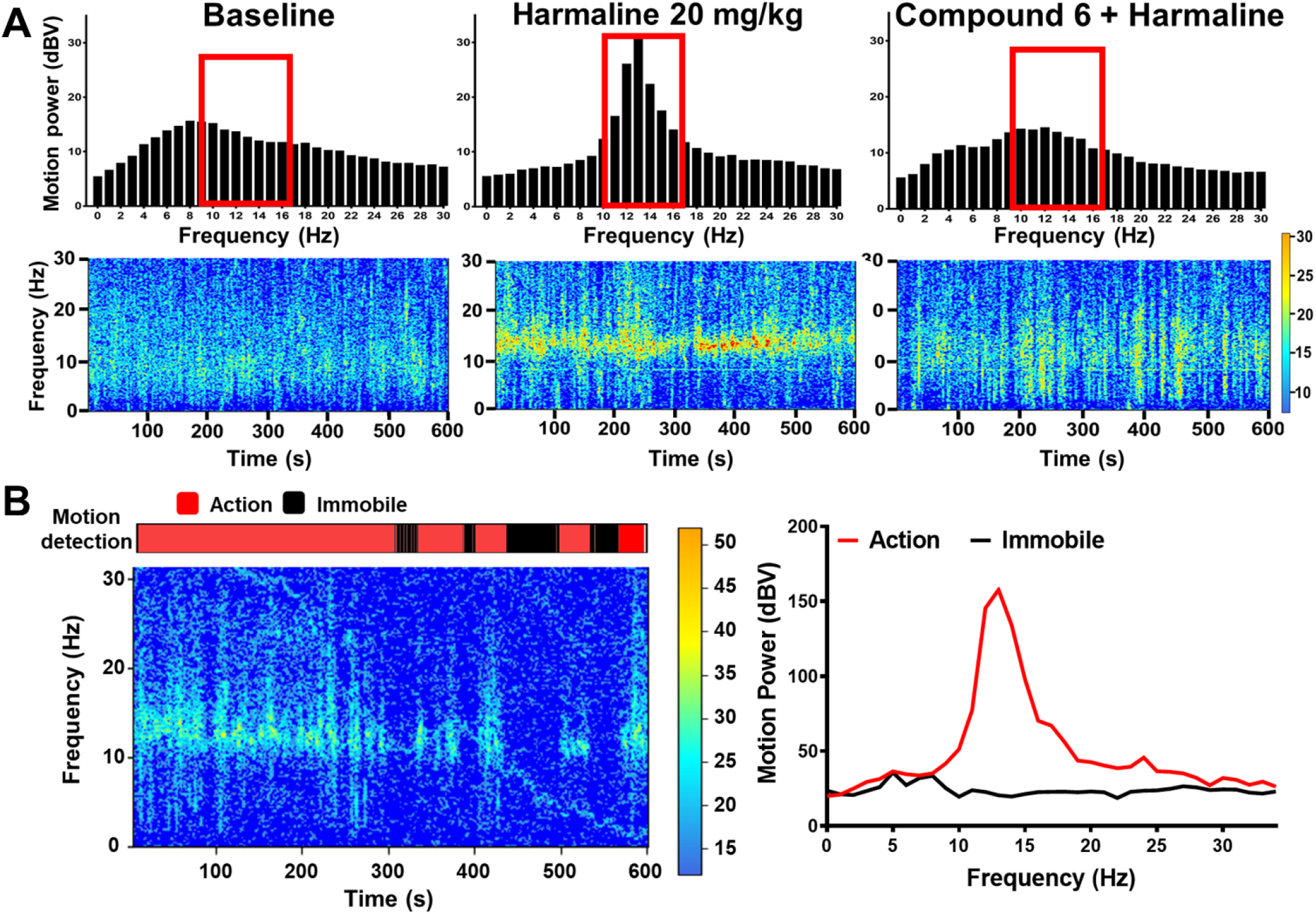
Representative figures of frequency domain-based motion power spectra and motion power spectrograms in mice treated with saline, harmaline and Compound 6 plus harmaline. **(A)** The motion waves in mice measured by a Tremor Monitor (San Diego Instruments, CA, USA) were converted into frequency domain motion power spectra (upper panels) and motion power heatmap spectrograms (lower panels) by a fast Fourier transformer. Note that harmaline induced tremor response at the frequency of 10-16 Hz in ICR mice, and Compound 6 attenuated harmaline-induced tremor activity. **(B)** A sample of 10-min motion power spectrograms (lower panel) and concurrent motor activity detected by SMART motion detector showing that the tremor activity occurs during movement (red region), but not immobile (black region) periods, suggesting that harmaline induces action tremor, but not rest tremor, in mice.

### Burrowing activity assessment

The burrowing activity of a mouse was measured as described previously (Ballon Romero *et al.*, 2020; Deacon, 2012) with modifications. For acclimatization, an empty tube was placed into the home cage, such that all five mice in the cage were exposed to the tube. On the day of testing, a separate cage was prepared for each mouse to be tested individually. The burrowing tube was filled with 200 g of food pellets. The burrowing activity was calculated as the weight of the expelled food pellet by subtracting the weight of the food pellet left at the end of the experiment from the initial weight (200g).

### Experimental protocol

As depicted in Fig. 3A, 10 min after harmaline (*s.c.*) injection, the mouse was placed in the tremor chamber and its tremor activity was measured for 10 min (pink shade). Right after the 10-min tremor measurement, the mouse was placed in a designated cage with the burrowing tube filled with 200 g of food pellets, and its burrowing activity was measured for another 10 min (grey shade). Then, the mouse was returned to the tremor chamber for tremor measurement. This tremor-burrowing measurement alternation was repeated for several cycles as indicated in each experiment. As harmaline-induced tremor is an action tremor, alternations were designed to refresh the alertness of the mouse once back in the tremor chamber, enhancing tremor consistency by replacing the rest period described in previous studies (Handforth *et al.*, 2018; Martin *et al.*, 2005) with the burrowing activity measurement.

**Fig. 3.**
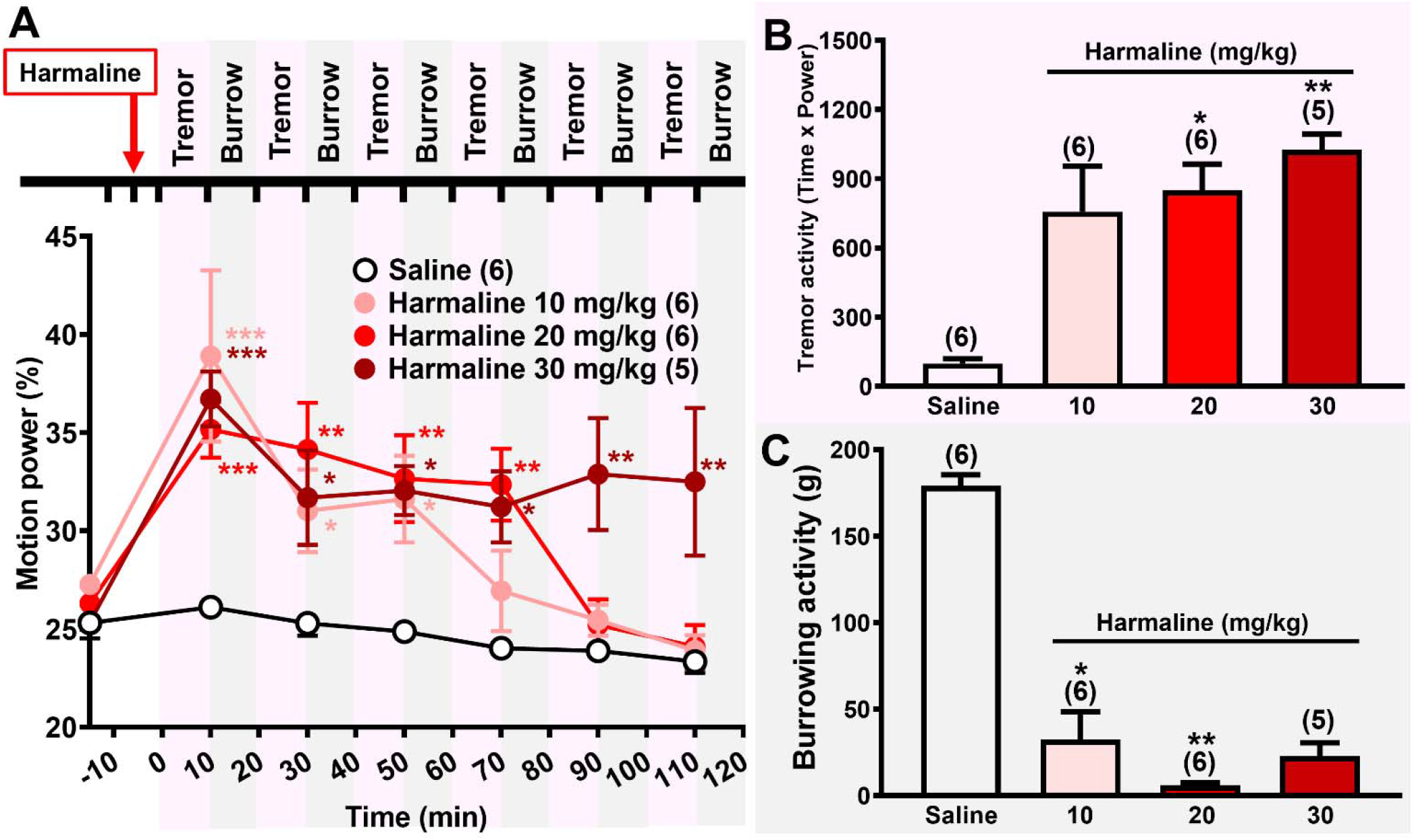
Dose-dependent effects of harmaline on tremor activity and burrowing activity in mice. **(A)** Time courses of the tremor activity induced by saline and 10, 20 and 30 mg/kg (*s.c.*) of harmaline. The tremor activity was measured for 10 min (pink shade), 15 min before, 5 min after, and then every 20 min after harmaline injection. Immediately after the tremor activity measurement, the mouse was placed in a chamber and its burrowing activity (grey shade) was measured for 10 min. Tremor activity and borrowing activity were measured alternatively for 120 min. The means of tremor activity for each 10-min period were plotted against time and compared among treatment groups. \**P* < 0.05, \*\**P* < 0.01, \*\*\**P* < 0.001 *vs.* Saline group, two-way ANOVA with repeated measures over time followed by Holm-Sidak’s *post hoc* test. **(B)** The total tremor activity, as measured by the area under curve (AUC) of the tremor activity against time, and **(C)** the burrowing activity, as measured by total displaced pellets, during 120 min-recording period in saline- or harmaline-treated mice. \**P* < 0.05, \*\**P* < 0.01 *vs.* Saline group, Kruskal-Wallis test followed by Dunn’s *post hoc* test. The numbers in the parentheses denoted the n number of mice tested in each group. Data are expressed as mean ± *S.E.M*.

### Drugs and chemicals

Compound 6, LAU 463, DK-I-56-1 and DK-I-58-1 were synthesized as previously described (Knutson *et al.*, 2018). Harmaline and propranolol were purchased from Sigma-Aldrich (St. Louis, MO, USA) and furosemide from Tocris Bioscience (Bristrol, UK). Compound 6, LAU463, DK-I-56-1, DK-I-58-1 (Fig. 1, A-D, respectively) and propranolol were dissolved in a vehicle containing 20% DMSO, 20% Cremophor^®^ EL (polyoxyethylene castor; Sigma-Aldrich) and 60% normal saline. Harmaline was dissolved in normal saline. Furosemide administered by *i.cb.* microinjection was dissolved in DMSO as reported previously (Chiou *et al.*, 2018; Liao *et al.*, 2016).

### Statistical analysis

Data were expressed as the mean ± S.E.M and the *n* number indicates the number of animals used. The two-way ANOVA followed by Holm-Sidak’s *post hoc* test was employed to examine the difference in the tremor intensity among different treatment groups across time. The one-way ANOVA followed by Holm-Sidak’s *post hoc* test was employed to examine the differences among treatment groups. However, if inhomogeneity was found in one-way ANOVA, as demonstrated in a significant Brown-Forsythe test, non-parametric Kruskal-Wallis followed by Dunn’s *post hoc* test was employed. Statistical differences were considered significant if *P* < 0.05.

## Results

### Harmaline consistently induced tremor and disrupted burrowing activity in mice

As depicted in Fig. 3A, harmaline at doses of 10, 20 and 30 mg/kg (*s.c.*), significantly induced tremor activity in mice as demonstrated by two-way ANOVA with repeated measures over time, which showed main effects of time [*F*(6,114) = 12.89, *P* < 0.001] and treatment [*F*(3,19) = 12.73, *P* < 0.001], and a significant interaction between time and treatment [*F*(18,114) = 2.778, *P* < 0.001]. The tremor activity peaked at 10-16 Hz as demonstrated in the motion power-frequency distribution plot and heatmap spectrogram (**Fig. 2A**). As shown in Fig 2B, tremor activity was detected during mobile (red) periods, but not during the immobile phase (black periods) of the harmaline-treated mice, suggesting that harmaline-induced tremor is an action tremor.

Harmaline-induced tremor activity reached peak at the first assessing time interval, 10-20 min after harmaline injection, and was similar among various dosage groups, but the time-dependent analysis with one-way ANOVA showed a significant difference among treatment groups [*F*(3,19) = 6.472, *P* = 0.003] (**Fig. 3A**). The tremor activity declined gradually and dose-dependently. The peak tremor activity lasted for at least 50, 75 and 115 min, respectively, induced by 10, 20 and 30 mg/kg harmaline. The area under curves (AUC) of the tremor activity over the 120 min-measuring period showed that the tremor-inducing effect of harmaline was dose-dependent (**Fig. 3B**). Besides inducing tremor, harmaline at doses of 10 and 20 mg/kg, significantly suppressed the burrowing activity in mice (**Fig. 3C**), but not at 30 mg/kg. The latter negative result might have been due to the lower n number of mice used in this experiment, or to unwanted detrimental effects induced by a high dose of harmaline since it is a neurotoxin that can act at many ion channels and neurotransmitter receptors (Du *et al.*, 1997; Jiang *et al.*, 2019; Zhan and Graf, 2012). Thus, harmaline at the dose of 20 mg/kg (s.c.) was chosen to induce tremor and burrowing activity deficit for subsequent pharmacological studies, with the measuring period limited to the first 75 mins after harmaline injection.

### Compound 6 (i.p.) suppressed tremor and restored burrowing activity in harmaline-treated mice

Similar to the protocol described in Fig. 3A, vehicle, Compound 6 (3 or 10 mg/kg) or propranolol (20 mg/kg) were *i.p.* administered 5 min prior to harmaline (20 mg/kg, *s.c.*) injection (**Fig. 4A**). Two-way ANOVA with repeated measurements over time showed main effects of time [*F*(4,164) = 41.59, *P* < 0.001] and treatment [*F*(3,41) = 12.57, *P* < 0.001], and a significant interaction between time and treatment [*F*(12,164) = 8.327, *P* < 0.001]. Both doses (3 and 10 mg/kg, *i.p.*) of Compound 6 and propranolol (20 mg/kg) significantly reduced tremor activity, as compared with the vehicle-treated group (**Fig. 4A**). The AUC derived from the time-dependent analysis showed that both doses of Compound 6 exerted significant anti-tremor effect, but propranolol showed the highest suppressive effect (**Fig. 4B**). One-way ANOVA showed a significant difference among treatment groups [*F*(3,41) = 12.85, *P* < 0.001]. However, one-way ANOVA followed by Holm-Sidak *post hoc* test demonstrated that the tremor-suppressant dose of propranolol failed to significantly restore the burrowing activity in harmaline-induced mice (*P* = 0.18), in contrast to Compound 6 at 3 mg/kg (*P* = 0.05) and 10 mg/kg (*P* = 0.05).

**Fig. 4.**
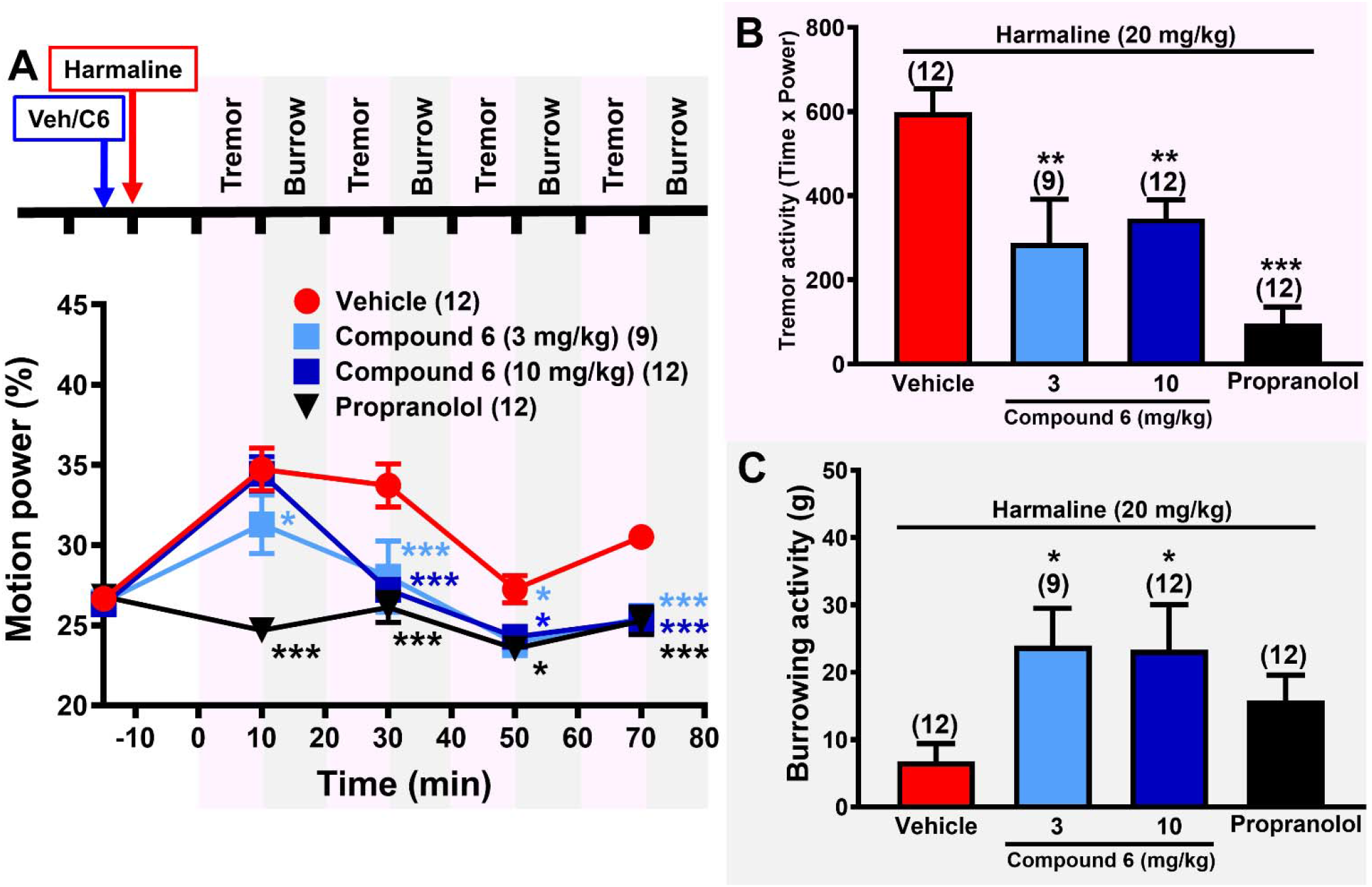
Effects of Compound 6, an α6GABA_A_R-selective PAM, and propranolol on harmalineinduced tremor and burrowing impairment in mice. **(A)** Time courses of effects of 3 and 10 mg/kg (*i.p.*) of Compound 6 and its vehicle, and propranolol (20 mg/kg) on harmaline (20 mg/kg, *s.c.*)-induced tremor. The tremor activity and burrowing activity were measured alternatively for 80 min as described in **Fig. 3A**. Compound 6 and vehicle were injected (*i.p.*) 5 min before harmaline treatment. \**P* < 0.05, \*\*\**P* < 0.001 *vs.* Saline group, two-way ANOVA with repeated measures over time followed by Holm-Sidak’s *post hoc* test. The total tremor activity **(B)** and burrowing activity **(C)** were measured as described in Figure 3 in harmaline-treated mice in various treatment groups. \**P* < 0.05, \*\**P* < 0.01, \*\*\**P* < 0.001 *vs.* Vehicle group, one-way ANOVA followed by Holm- Sidak’s *post hoc* test. The numbers in the parentheses denoted the n number of mice tested in each group. Data are expressed as mean ± *S.E.M*. Veh: vehicle, C6: Compound 6.

### Furosemide (i.cb.) antagonized effects of Compound 6 (i.p.) on tremor and burrowing activities in harmaline-treated mice

In order to discern whether cerebellar α6GABA_A_Rs are involved in the anti-tremor effect of Compound 6, we treated mice with co-administration of *i.cb.* furosemide (10 nmol) and *i.p.* Compound 6 (10 mg/kg), 5 min before harmaline injection (**Fig. 5A**). Two-way ANOVA with repeated measures over time showed main effects of time [*F*(4,164) = 41.59, *P* < 0.001] and treatment [*F*(3,41) = 12.57, *P* < 0.001], and a significant interaction between time and treatment [*F*(12,164) = 8.327, *P* < 0.001] (**Fig. 5A**). In both time-dependent analyses (yellow diamonds, **Fig. 5A**) and the derived AUC of motion power (yellow bar, **Fig. 5B**) in harmaline-treated mice, *i.cb.* furosemide alone had no effect on the tremor activity. Co-administered of Compound 6 with *i.cb.* microinjection of furosemide (green bar, **Fig. 5B**), but not its vehicle (blue bar, **Fig. 5A**), did not display significant anti-tremor effect, suggesting *i.cb.* furosemide, at the dose without affecting tremor activity *per se*, significantly antagonizes the anti-tremor effect of Compound 6. Interestingly, the restorative effect of Compound 6 on burrowing-activity in harmaline-treated mice was also completely reversed by *i.cb.* furosemide (green bar, **Fig. 5C**) while *i.cb.* furosemide *per se* did not affect the burrowing activity (yellow bar, **Fig. 5C**).

**Fig. 5.**
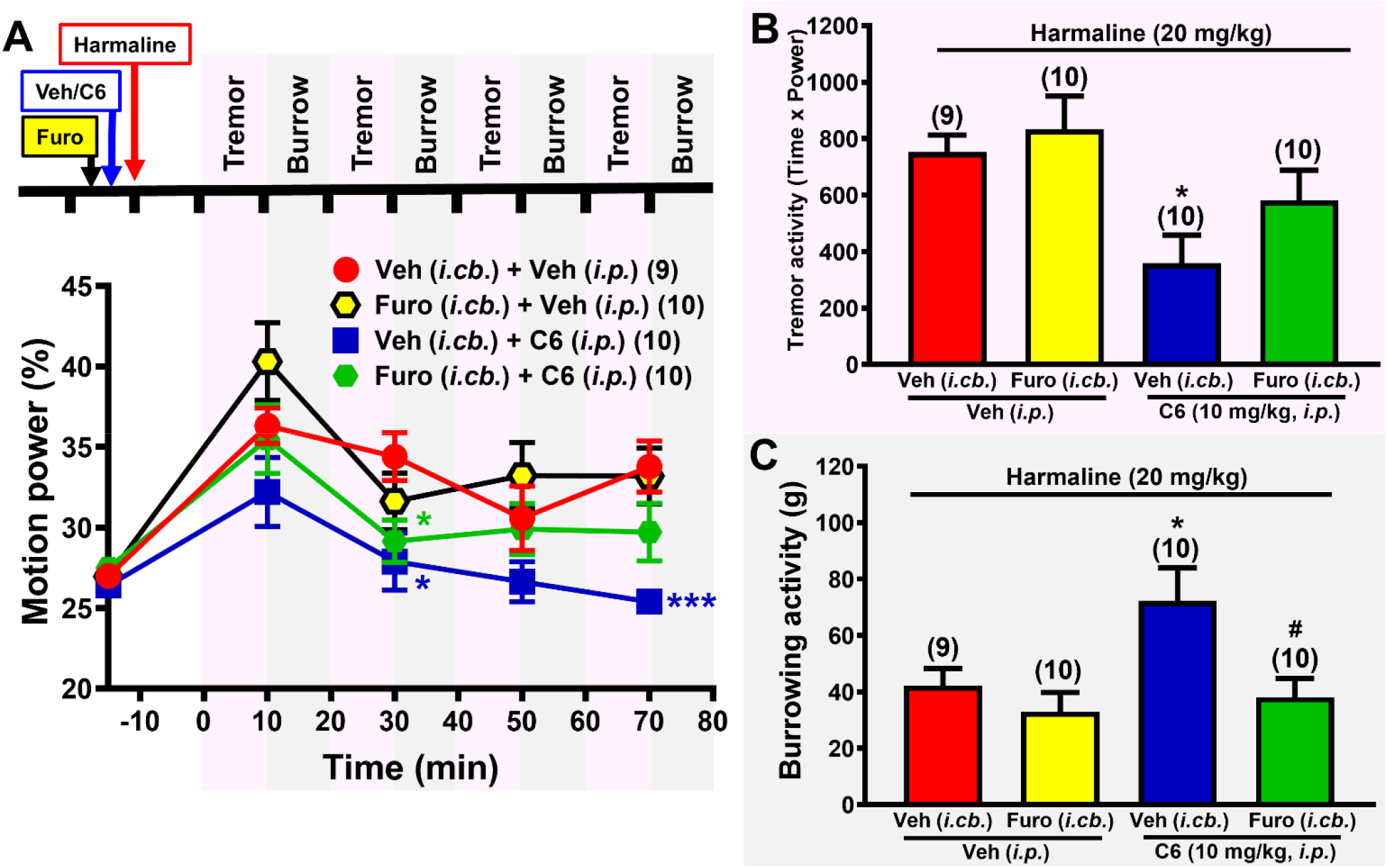
Effects of intra-cerebellar microinjection (*i.cb.*) of furosemide, an α6GABA_A_R antagonist, on anti-tremor and burrow-restorative effects of Compound 6 in harmaline-treated mice. **(A)** Time courses of effects of 10 mg/kg (*i.p.*) of Compound 6 and its vehicle without and with *i.cb.* furosemide (10 nmol) co-treatment on the tremor activity of harmaline-treated mice. Tremor activity and burrowing activity were measured alternatively for 80 min as described in **Fig. 4A**. \**P* < 0.05, \*\*\**P* < 0.001 *vs.* Saline group, two-way ANOVA with repeated measures over time followed by Holm-Sidak’s *post hoc* test. The total tremor activity **(B)** and burrowing activity **(C)** were measured as described in Figure 3 in harmaline-treated mice pretreated with Compound 6 alone or in combination with *i.cb.* furosemide. \**P* < 0.05, \*\**P* < 0.01, \*\*\**P* < 0.001 *vs.* Vehicle group, ^#^*P* < 0.05 *vs.* Veh (i.cb.)+C6 group, one-way ANOVA followed by Holm-Sidak’s *post hoc* test. Furo: furosemide, Veh: vehicle, C6: Compound 6.

### Compound 6 analogues and deuterated derivatives displayed similar anti-tremor and burrowing-restorative effects in harmaline-treated mice

Compound 6 and its structural analogue, LAU463, as well as their respective deuterated derivatives, DK-I-56-1 and DK-I-58-1, were previously demonstrated to possess similar selectivity towards α6GABA_A_R as PAMs. In the present study, administration of DK-I-56-1 (half-filled square symbols, **Fig. 6A**), LAU-463 (filled diamonds, **Fig. 6B**) and DK-I-58-1 (half-filled diamonds, **Fig. 6C**) at the of 3 and 10 mg/kg (*i.p.*) demonstrated time and treatment-dependent suppression of harmaline-induced tremor in mice, lasting for almost 1 hr. By comparing the AUC derived from time-dependent analysis showed that each drug group significantly suppressed tremor activity as compared with vehicle group, except DK-I-58-1 at the dose of 3 mg/kg. Interestingly, as compared with the vehicle group within each drug group, all treatment groups significantly restored burrowing activity in harmaline-treated mice (**Fig. 6E**), similar to the effect of Compound 6, but not of propranolol (**Fig. 4C**),

**Fig. 6.**
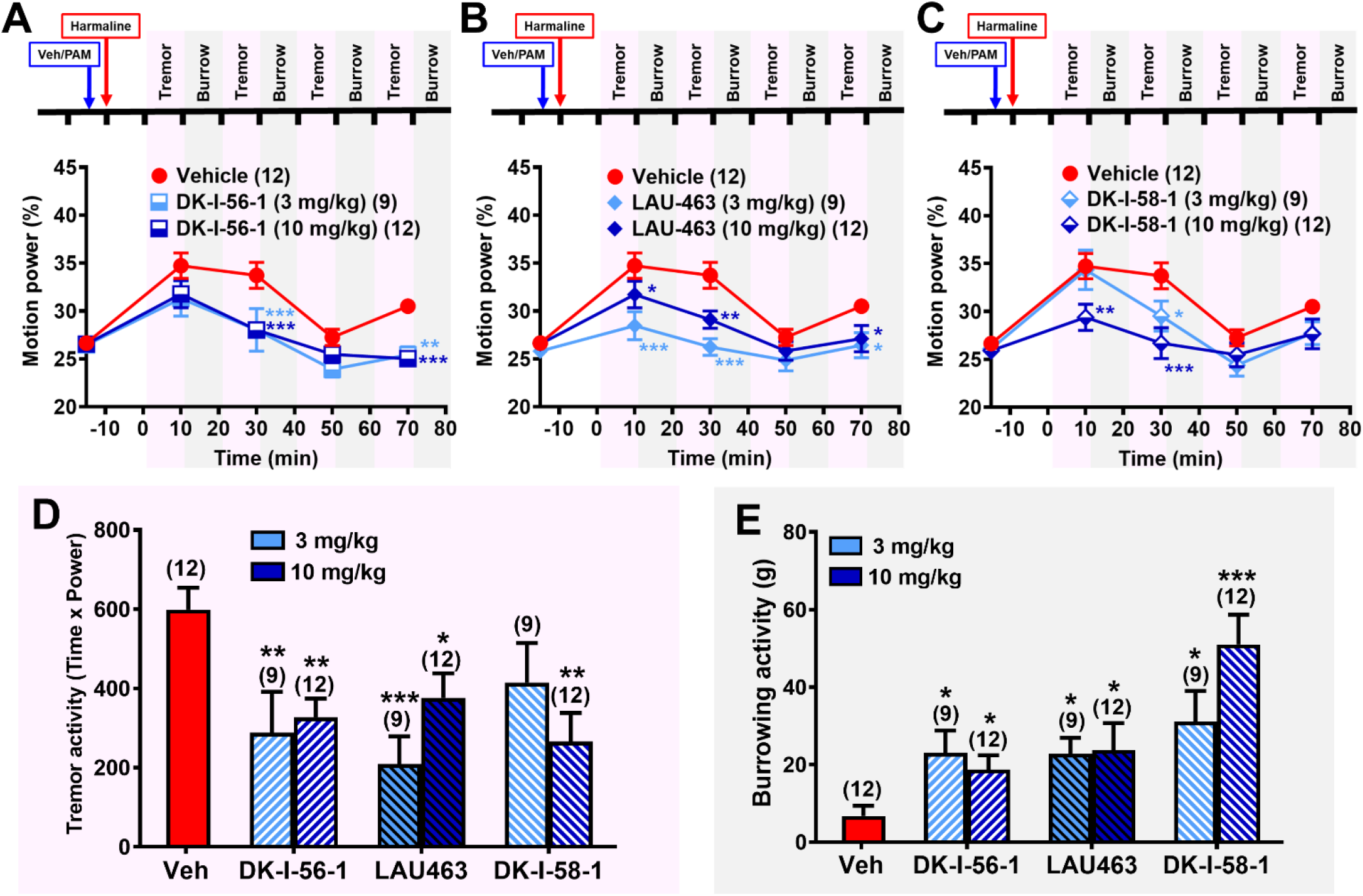
Effects of DK-I-56-1, LAU463 and DK-I-58-1, structural analogues of Compound 6, on harmaline-treated mice. LAU463 is a structural analogue of Compound 6. DK-I-58-1 and DK-I-56-1 are their respective deuterated derivatives. These compounds are all α6GABA_A_R PAMs with similar α6GABA_A_R-selectivity and efficacy while deuterated derivatives have longer half-lives. Similar procedure as described in Fig. 4A were performed with Compound 6 replaced with **(A)** DK-I-56-1 (half-filled square symbols), **(B)** LAU463 (diamond symbols) or **(C)** DK-I-58-1 (half-filled diamond symbols), at doses of 3 and 10 mg/kg (*i.p.*). \**P* < 0.05, \*\**P* < 0.01, \*\*\**P* < 0.001 *vs.* Saline group, two-way ANOVA with repeated measures over time followed by Holm-Sidak’s *post hoc* test. The total tremor activity **(D)** and burrowing activity **(E)** were measured alternatively for 80 min as described in Figure 3 in various treatment groups. \**P* < 0.05, \*\**P* < 0.01, \*\*\**P* < 0.001 *vs.* Vehicle group, one-way ANOVA followed by Holm-Sidak’s *post hoc* test. Veh: vehicle, PAM: positive allosteric modulator.

## Discussion

### α6GABA_A_R-selective PQ compounds suppressed tremor and restored burrowing activity in harmaline-treated mice

In this study, we found that harmaline at 10-30 mg/kg (*s.c.*) induced significant action tremor at 10-16 Hz, with a similar maximal tremor activity but a dose-dependent action duration in ICR mice (**Fig. 3**), consistent with a previous report (Martin *et al.*, 2005). The predictive validity of harmaline-induced tremor in modeling ET in patients was strongly supported by the finding that it responded to clinically used ET-relieving agents, such as propranolol, primidone, alcohol, benzodiazepines, gabapentin, gammahydroxybutyrate, 1-octanol, and zonisamide, and was exacerbated by drugs that worsen ET, such as tricyclics and caffeine (Handforth, 2012). In this action tremor model, we provided a proof-of-concept pre-clinical study indicating that α6GABA_A_Rselective PAMs are a novel pharmacotherapy for ET.

Compound 6 as well as its three structural analogues effectively suppressed tremor activity and restored burrowing activity in harmaline-treated mice. Whereas Compound 6 and LAU-463 are structural analogues with different functional groups on the A-ring, DK-I-56-1 and DK-I-58-1 are their respective derivatives with a deuterated methoxy group on the D-ring (**Fig. 1**). These deuterated derivatives, compared with their parent compounds, had a similar potency and the same α6GABA_A_R-selectivity in enhancing GABA currents of recombinant α6βγGABA_A_Rs expressed in Xenopus oocytes (Knutson *et al.*, 2018; Treven *et al.*, 2018) but displayed longer half-lives in rodents (Knutson *et al.*, 2018). Although the merit of longer half-lives of deuterated compounds, cannot be observed in these behavioral assays that only lasted for 90 minutes, for future research and a possible clinical development of α6GABA_A_R-selective PQ compounds, it was important to also investigate the activity of these deuterated compounds in these assays.

The burrowing activity in laboratory rodents is one of their “Activities of Daily Living” (ADL) (Deacon, 2012), and has been used as an indicator of well-being (Jirkof, 2014), because it is negatively associated with stress (Van Loo *et al.*, 2007) or chronic pain (Huang *et al.*, 2013) in rodents. In addition, a deficit of the burrowing activity has also been reported in mouse models of neurodegenerative diseases, such as Alzheimer’s disease (Geiszler *et al.*, 2016) and Parkinson’s disease (Baumann *et al.*, 2016). The finding that harmaline markedly disrupted the burrowing activity in mice (**Fig. 3C**) suggests that this ET animal model mimics not only the action tremor but also the reduced ADL score manifested in ET patients because of the motor disorder-induced distress that severely affects their well-being. Interestingly, α6GABA_A_R-selective PQ compounds also restored the burrowing activity in harmaline-treated mice (**Fig. 4C** and **Fig. 6E**) in addition to attenuating their tremor activity. In contrast, propranolol, the first-line medication for essential tremor, did not restore the burrowing activity in mice (**Fig. 2D**). This is probably due to the reduction of the locomotor activity induced by propranolol (Poli and Palermo-Neto, 1985), which also causes muscle weakness in humans (Deuschl *et al.*, 2011). Since α6GABA_A_R PAMs can both reduce tremor activity and restore burrowing activity in harmaline-treated mice, they may be viable candidates to complement current anti-tremor drugs.

### Cerebellar α6GABA_A_Rs are the site of action of anti-tremor effects of α6GABA_A_R-selective PQ compounds

The finding that *i.cb.* microinjection of furosemide, an α6GABA_A_R-selective antagonist (Korpi and Luddens, 1997), antagonized the anti-tremor effect of Compound 6 (**Fig. 3, B & C**) suggests that the α6GABA_A_Rs in the cerebellum are the site of action of Compound 6. A previous study indicated that gaboxadol, a selective agonist of the δ-subunit containing GABA_A_Rs (δGABA_A_Rs), including the α6βδ subtype of GABA_A_Rs, significantly suppressed harmaline-induced tremor in wild type, but not in *Gabra6*^-/-^or *Gabrd*^-/-^mice (Handforth *et al.*, 2018), suggesting that the α6βδGABA_A_R subtype is involved in harmaline-induced tremor. Our previous electrophysiological study on oocyte expressing various recombinant GABA_A_R subtypes demonstrated that Compound 6 exerted its PAM activity at both α6βγ2 and α6βδ containing α6GABA_A_Rs although having a lower efficacy for the α6βδ subtype (Chiou *et al.*, 2018). Even though, it enhanced the GABA current of α6βδGABA_A_Rs to 270% at 3 μM (Chiou *et al.*, 2018). Thus, it is likely that cerebellar α6βδGABA_A_Rs are the action target of Compound 6. On the other hand, a contribution of the α6βγ2GABA_A_R subtype in the anti-tremor effect of Compound 6 cannot be excluded. Compound 6 could reduce the neuronal excitability of granule cells by enhancing GABAergic transmission at extrasynaptic α6βδGABA_A_Rs as well as at α6βγ2GABA_A_Rs at Golgi cell-granule cell synapses (Nusser *et al.*, 1998), thus ultimately reducing the excitatory input of granule cells and hence the mis-firing of PCs.

### Current essential tremor therapy is insufficient

Currently, essential tremor is symptomatically treated with medicines that can suppress tremor activity, like ß-blockers, anti-epileptics, neuron stabilizers and GABA-related drugs (Gironell, 2014; Shanker, 2019). Among these, propranolol and primidone remain the first line medications for relieving tremor in patients with essential tremor, even decades after their initial applications (Shanker, 2019). A combination of these two drugs yields a better response than using either of them alone (Zesiewicz *et al.*, 2005). However, one third of patients developed drug resistance (Deuschl *et al.*, 2011), and a significant proportion of the patients are even refractory to both medications (Louis *et al.*, 2010). The reported adverse effects differ between these two drugs with acute reactions more common with primidone and chronic reactions with propranolol (Bain, 1997). The acute side effects associated with primidone are sedation, nausea and vertigo. They are manifest after the first dose and tend to subside on chronic administrations (Koller and Royse, 1986; Sasso *et al.*, 1990). On the other hand, chronic propranolol may lead to bradycardia, hypotension and breathlessness, especially when higher doses are used. Therefore, propranolol is contraindicated in conditions such as chronic obstructive pulmonary disease, asthma, severe peripheral vascular disease and diabetes (Winkler and Young, 1974).

Limited data from randomized controlled trials are available to support the use of other medications in essential tremor, including topiramate, alprazolam, gabapentin, and other betablockers besides propranolol (e.g., atenolol, nadolol, and sotalol) (Deuschl *et al.*, 2011; Haubenberger and Hallett, 2018; Schaefer *et al.*, 2018; Zesiewicz *et al.*, 2011). Randomized controlled trials have shown no significant benefit for several other drugs for essential tremor, including levetiracetam, amifampridine, flunarizine, trazodone, pindolol, acetazolamide, mirtazapine, nifedipine, and verapamil (Zesiewicz *et al.*, 2011). As a result, the need for an effective anti-tremor drug that is well-tolerated is urgent and obvious.

### α6GABA_A_R-selective PQ compounds as a novel therapy for essential tremor

We have previously demonstrated that Compound 6, one of the first α6GABA_A_R-selective PAMs identified (Knutson *et al.*, 2018; Varagic *et al.*, 2013), via acting at cerebellar α6GABA_A_Rs, ameliorated disrupted prepulse inhibition (PPI) of the startle response (Chiou *et al.*, 2018), social withdrawal and cognitive impairment (Chiou *et al.*, 2019) in animal models mimicking schizophrenia. By acting via α6GABA_A_Rs at trigeminal ganglia, it inhibited nociceptive activation of the trigeminovascular system and migraine-like grimaces in animal models of migraine (Fan *et al.*, 2018; Tzeng *et al.*, 2020). And, a deuterated Compound 6 analogue, DK-I-56-1, prevented and reduced trigeminal neuropathic pain in a rat model (Vasovic *et al.*, 2019). Here, we further substantiated the effectiveness of α6GABA_A_R-selective PQ compounds in an animal model of essential tremor. The effective dose range (3-10 mg/kg, *i.p.*) of Compound 6 in suppressing tremor was similar to that in mouse models of schizophrenia (Chiou *et al.*, 2018; Chiou *et al.*, 2019) and migraine (Tzeng *et al.*, 2020). At these doses, Compound 6 did not impair motor functions in rats (Knutson *et al.*, 2018), nor display sedative (Wu and Chiou, 2016) or hypolocomotor activity (Chiou *et al.*, 2019) in mice. In contrast to the GABA binding site agonist, gaboxadol (THIP), which directly activates all α4βδ and α6βδ GABA_A_Rs in nervous tissues (Brown *et al.*, 2020; Meera *et al.*, 2011) and exhibits sedation and motor-incoordination (Wafford and Ebert, 2006), α6GABA_A_R-selective PAMs can only allosterically modulate the activity of those α6βδ and α6βγ2 GABA_A_Rs, which are activated by GABA in a certain task, explaining the absence of such side effects. In contrast to BDZs and gaboxadol, α6GABA_A_R-selective PQ compounds may thus exert therapeutic effects without sedative or motor-impairing activity.

In addition, the investigated α6GABA_A_R-selective PQ compounds have a satisfactory safety pharmacology profile. In a panel of radioligand binding assays targeting 46 CNS receptors/channels/enzymes, these PQ compounds displayed an affinity for GABA_A_R binding that was more than 1000-fold higher than that for 45 other off targets, including the hERG channel, an arrhythmia risk potassium channel (Knutson *et al.*, 2018). In cytotoxicity tests on HEK293 (kidney) and HEPGS (liver) cells, PQ compounds at the concentration up to 400 μM did not display significant cytotoxicity (Knutson *et al.*, 2018). However, due to their α6GABA_A_R selectivity, these PQ compounds only modulated the excitatory input of mossy fibers, but not that of climbing fibers in cerebellum, possibly explaining their incomplete suppression of the tremor activity. Nevertheless, in view of their novel target-acting property, satisfactory safety pharmacology profile, and absence of side effects, the investigated α6GABA_A_R-selective PAMs can be developed as an alternative monotherapy or a non-toxic add-on therapy for the treatment of ET.

## Acknowledgement

This study was supported by grants from the Ministry of Science and Technology, Taiwan (MOST 104-2923-B-002-006-MY3, MOST 108-2320-B-002-029-MY3 and 109-2320-B-002-042-MY3 to LCC), National Health Research Institutes, Taiwan (NHRI-EX107-10733NI to LCC), Ministry of Education, Taiwan (107M4022-3 to LCC), UCSI University Research Excellence and Innovation Grant (REIG-FPS-2020/065 to MTL) National Institutes of Health, USA (R01AA029023 and R01MH096463 to JC), and the National Science Foundation, USA, Division of Chemistry (CHE-1625735 to JC). We thank the Milwaukee Institute of Drug Discovery for the mass spectroscopy. LRW was supported by the Austrian Science Fund FWF grant W1232 (Graduate School Program MolTag).

## Authors’ Contributions

YHH conducted experiments, analyzed data, and wrote the paper; MTL and WS contributed to data interpretation, data analysis, result discussion, and paper writing; DEK, LRW, DS, MDM and JC synthesized and provided compounds; LCC provided supervision, designed experiments, analyzed data, and wrote the paper.

